## Supplementary for "Human-environment feedback and the consistency of proenvironmental behavior"

### Supplementary material for Human-environment feedback and the consistency of proenvironmental behavior

In this supplementary material, we expound the mathematical results stated in the main text. We define a piecewise deterministic Markov process (PDMP) coupling environmental and behavioral dynamics. Taking the limit when the population size tends to infinity leads to a deterministic two-dimensional dynamical system. We study the stability of this system and perform the bifurcation analysis. We also study the fluctuations of the PDMP around the deterministic limit. Finally, we provide details about the simulation algorithm of the stochastic population-environment process.

†Co-last authors

### 1 Properties of the stochastic system

In the main text [3], we introduced a time continuous two-dimensional stochastic process describing the joint dynamics of the stochastic populational level and the macroscopic environment. The individuals are characterized by a parameter $u \in \{A, B\}$ describing two possible behaviors: active (A) or baseline (B). The process $((X_t^N, E_t^N), t \geq 0)$ is a PDMP scaled by the fixed population size $N$. The perceived environmental state at time t is quantified by $E_t^N$ which takes values in the compact interval $[l_A, l_B]$, $l_A < l_B$. The deterministic dynamics of the environmental state are shaped by the function $h$ defined in [3]-Equation (1) as

$$h(x, e) = \ell e(l_A x + l_B(1 - x) - e) \tag{1}$$

$(x, e)$ being the joint population-environment state. The frequency dynamics of the active individual behavior are given by the pure jump process $(X_t^N, t \geq 0)$ whose transitions are driven by two possible switches due either to the social or tto the environmental influence. An individual with behavior $u \in \{A, B\}$ can switch to $v \neq u \in \{A, B\}$ at rate $\kappa N \lambda_v^N(x)$ in response to the social influence and $\tau_v(e)$ (defined in [3]-Equation (3) respectively by $\tau_A(e) = \tau(e - l_A)$ and $\tau_B(e) = \tau(l_B - e)$) in response to the environmental influence.

Using the classical pathwise representation of the PDMP driven by a Poisson point measure, we easily obtain the semi-martingale decomposition of $(X^N, E^N)$ for fixed N, as

$$\begin{cases} X_t^N = X_0^N + \int_0^t P^N(X_s^N, E_s^N) ds + M_t^N \\ E_t^N = E_0^N + \int_0^t h(X_s^N, E_s^N) ds, \end{cases} \tag{2}$$

where $P^N$ is given by

$$P^N(x, e) = \kappa x(1 - x) N[\lambda_A^N(x) - \lambda_B^N(x)] + (1 - x)\tau_A(e) - x\tau_B(e) \tag{3}$$

and $M_t^N$ is a square-integrable martingale whose quadratic variation is

$$\begin{aligned} \langle M^N \rangle_t = \; & \frac{1}{N} \int_0^t \kappa X_s^N (1 - X_s^N) N[\lambda_A^N(X_s^N) + \lambda_B^N(X_s^N)] ds \\ & + \frac{1}{N} \int_0^t \tau_A(E_s^N)(1 - X_s^N) + \tau_B(E_s^N) X_s^N ds. \end{aligned} \tag{4}$$

### 2 Convergence to the solution of a system of ordinary differential equations

The next theorem describes the limiting behavior of the process $(X_t^N, E_t^N)_{t \geq 0}$ when $N$ goes to infinity.

We assume that there exist positive constants $\gamma_A$, $\gamma_B$, $\delta_A$ and $\delta_B$ such that for all $x \in [0, 1]$,

$$
\begin{aligned}
N\lambda_A^N(x) &\underset{N\to+\infty}{\longrightarrow} \lambda_A(x) = \gamma_A + \delta_A x,\\
N\lambda_B^N(x) &\underset{N\to+\infty}{\longrightarrow} \lambda_B(x) = \gamma_B + \delta_B(1-x).
\end{aligned} \tag{5}
$$

**Theorem 1.** *When $N \to +\infty$, under Assumption* (5) *and if* $(X_0^N, E_0^N) \to (x_0, e_0) \in [0,1] \times [l_A, l_B]$, *then for all* $T > 0$, *the process* $((X_t^N, E_t^N), t \in [0,T])$ *converges in probability in* $\mathbb{D}([0,T], [0,1] \times [l_A, l_B])$ *to the unique solution* $((x_t, e_t), t \in [0,T])$ *of the following system of ordinary differential equations (ODEs)*

$$
\begin{aligned}
\frac{dx_t}{dt} &= \kappa x_t(1-x_t)[\gamma_A - \gamma_B + ((\delta_A+\delta_B)x_t - \delta_B)] + \tau(e_t - l_A - (l_B - l_A)x_t),\\
\frac{de_t}{dt} &= \ell e_t(l_A x_t + l_B(1-x_t) - e_t),
\end{aligned} \tag{6}
$$

*with initial conditions* $(x_0, e_0)$.

*Proof.* The proof is based on a tightness-uniqueness scheme, as in [1, section 3.1], see also [4].

The processes $((X_t^N, E_t^N), t \in [0,T])$ evolve in a compact set so they obviously satisfy for any $T > 0$,

$$
\sup_N \mathbb{E}\left(\sup_{t\in[0,T]} \|(X_t^N, E_t^N)\|^3\right) < +\infty. \tag{7}
$$

The tightness of their laws can be deduced from (2) and from this uniform moment estimate. The relative compactness of this family of laws in the set of probability measures $\mathbb{D}([0,T], [0,1] \times [l_A, l_B])$ follows, as well as the existence of a limiting value $Q$. To identify the latter, we define for $z \in \mathbb{D}([0,T], [0,1] \times [l_A, l_B])$ and for any $t > 0$ the function

$$
\Psi_t(z) = z_t - z_0 - \int_0^t H(z_s)ds,
$$

where $H = (P, h)$ and $P$ is the limit of $P^N$ under Assumption (5), given for all $(x, e) \in [0,1] \times [l_A, l_B]$ by

$$
P(x,e) = \kappa x(1-x)[\gamma_A - \gamma_B + ((\delta_A+\delta_B)x - \delta_B)] + \tau(e - l_A - (l_B - l_A)x).
$$

The assumptions yield $|\Psi_t(z)| \le C \sup_{t\le T}(1 + (z_t))$. Then by Eq. (7), the function $\Psi_t$ is uniformly integrable and

$$
\mathbb{E}_Q(|\Psi_t(z)|) = \lim_N \mathbb{E}(|\Psi_t(Z^N)|) = \lim_N \mathbb{E}(|\Psi_t^N(Z^N)|) = \lim_N \mathbb{E}(|M_t^N|),
$$

where $\psi^N$ is defined as $\psi$ with $P^N$ instead of $P$. Using $\lim_N \mathbb{E}(|M_t^N|) \le \lim_N \left(\mathbb{E}\left(|M_t^N|^2\right)\right)^{\frac{1}{2}}$ and Eq. (4), we obtain that $\lim_N |M_t^N| = 0$ a.s. Then under $Q$, the limiting process is solution of $z_t - z_0 - \int_0^t H(z_s)ds = 0$, i.e. is solution of (6). The uniqueness follows from the Cauchy-Lipschitz Theorem. We finally conclude that the processes $((X_t^N, E_t^N), t \in [0,T])$ converge uniformly in law (and then in probability) for any time $t \in [0,T]$ to the unique solution of the dynamical system (6).

□

### 3 Properties of the dynamical system (6)

We are interested in the long-term behavior of the dynamical system (6). Without loss of generality, we simplify the dynamical system by introducing a new parameterization.

#### 3.1 Parameterization of the model

We choose the following parameterization. We introduce new perceived environment and time variables $v$ and $\tilde{\tau}$ by setting $v = \dfrac{e - l_A}{l_B - l_A}$ and $\tilde{\tau} = \tau(l_B - l_A)t$ and we rescale $r = \dfrac{\ell l_B}{\tau(l_B - l_A)}$. We also define $a = \frac{\kappa(\gamma_A - \gamma_B - \delta_B)}{\tau(l_B - l_A)}$, $b = \frac{\kappa(\delta_A + \delta_B)}{\tau(l_B - l_A)}$ and $c = \frac{(l_B - l_A)}{l_B}$. In the new parameterization, system (6) becomes

$$\begin{aligned} \frac{dx_{\tilde{\tau}}}{d\tilde{\tau}} &= -f(x_{\tilde{\tau}}) + v_{\tilde{\tau}}, \\ \frac{dv_{\tilde{\tau}}}{d\tilde{\tau}} &= r(cv_{\tilde{\tau}} + (1-c))(1 - x_{\tilde{\tau}} - v_{\tilde{\tau}}), \end{aligned} \tag{8}$$

with $\forall \tilde{\tau} \geq 0,\ (x_{\tilde{\tau}}, v_{\tilde{\tau}}) \in [0,1]^2$ and

$$f(x) = x((x-1)(a+bx)+1). \tag{9}$$

Since we assumed that $\kappa,\ \tau,\ \ell,\ \gamma_A,\ \gamma_B,\ \delta_A,\ \delta_B$ and $l_B > l_A > 0$ are positive, it follows that $c \in ]0,1]$ and b and r are positive.

*Proof.* Using the parametrisation, we have:

$$\frac{dv_{\tilde{\tau}}}{de_{\tilde{\tau}}} = \frac{1}{l_B - l_A} \text{ and } \frac{dt}{d\tilde{\tau}} = \frac{1}{\tau(l_B - l_A)}.$$

Thus, we have for the first equation

$$\begin{aligned} \frac{dx_{\tilde{\tau}}}{d\tilde{\tau}} &= \frac{dx_{\tilde{\tau}}}{dt}\frac{dt}{d\tilde{\tau}} \\ &= \frac{\kappa x_{\tilde{\tau}}(1-x_{\tilde{\tau}})[\gamma_A - \gamma_B + ((\delta_A + \delta_B)x_{\tilde{\tau}} - \delta_B)] + \tau(e_{\tilde{\tau}} - l_A - (l_B - l_A)x_{\tilde{\tau}})}{\tau(l_B - l_A)} \\ &= x_{\tilde{\tau}}(1 - x_{\tilde{\tau}})\left[\frac{\kappa(\gamma_A - \gamma_B - \delta_B)}{\tau(l_B - l_A)} + \frac{\kappa(\delta_A + \delta_B)}{\tau(l_B - l_A)}x_{\tilde{\tau}}\right] + \frac{e_{\tilde{\tau}} - l_A}{l_B - l_A} - x_{\tilde{\tau}} \\ &= -x_{\tilde{\tau}}((x_{\tilde{\tau}} - 1)(a + bx_{\tilde{\tau}}) + 1) + v_{\tilde{\tau}}. \end{aligned}$$

For the second equation, we have :

$$
\begin{aligned}
\frac{dv_{\tilde{\tau}}}{d\tilde{\tau}} &= \frac{dv_{\tilde{\tau}}}{de_{\tilde{\tau}}}\,\frac{de_{\tilde{\tau}}}{dt}\,\frac{dt}{d\tilde{\tau}} \\
&= \frac{1}{l_B - l_A}\ell e_{\tilde{\tau}}(l_A x_{\tilde{\tau}} + l_B(1 - x_{\tilde{\tau}}) - e_{\tilde{\tau}})\frac{1}{\tau(l_B - l_A)} \\
&= \frac{\ell}{\tau}\left(\frac{e_{\tilde{\tau}} - l_A + l_A}{l_B - l_A}\right)(l_A x_{\tilde{\tau}} + l_B(1 - x_{\tilde{\tau}}) - e_{\tilde{\tau}} + l_A - l_A)\,\frac{1}{l_B - l_A} \\
&= \frac{\ell}{\tau}\left(v_{\tilde{\tau}} + \frac{l_A}{l_B - l_A}\right)(l_A(x_{\tilde{\tau}} - 1) + l_B(1 - x_{\tilde{\tau}}) - e_{\tilde{\tau}} + l_A)\,\frac{1}{l_B - l_A} \\
&= \frac{\ell l_B}{\tau(l_B - l_A)}\left(v_{\tilde{\tau}} + \frac{l_B}{l_B - l_A} - 1\right)(1 - x_{\tilde{\tau}} - v_{\tilde{\tau}}) \\
&= r\,(c v_{\tilde{\tau}} + 1 - c)\,(1 - x_{\tilde{\tau}} - v_{\tilde{\tau}})\,.
\end{aligned}
$$

□

Let us now study the fixed points and the possible bifurcations of System (8).

#### 3.2 Stability analysis

We recall that an equilibrium $(x^*, v^*)$ of System (8) is a point that satisfies $\frac{dx_{\tilde{\tau}}}{d\tilde{\tau}}(x^*, v^*) = \frac{dv_{\tilde{\tau}}}{d\tilde{\tau}}(x^*, v^*) = 0$. Thus, the equilibria lie at the intersection of the cubic $v = f(x)$ and the line $v = 1 - x$. To study this intersection, we introduce the auxiliary function g given by

$$g(x) = f(x) + x - 1, \ \forall x \in [0, 1]. \tag{10}$$

**Proposition 2.** *System* (8) *has 1, 2 or 3 stationary points.*

*Proof.* The equilibria of System (8) belong to the intersection between the cubic $v = f(x)$ and the line $v = 1 - x$ in the unit square. We are then looking at the zeros of the function $g$ on $[0, 1]$. The function $g$ is a polynomial of degree 3 and admits at most 3 roots. The system has at least one equilibrium on [0,1] because $g(0) = -1$ and $g(1) = 1$. □

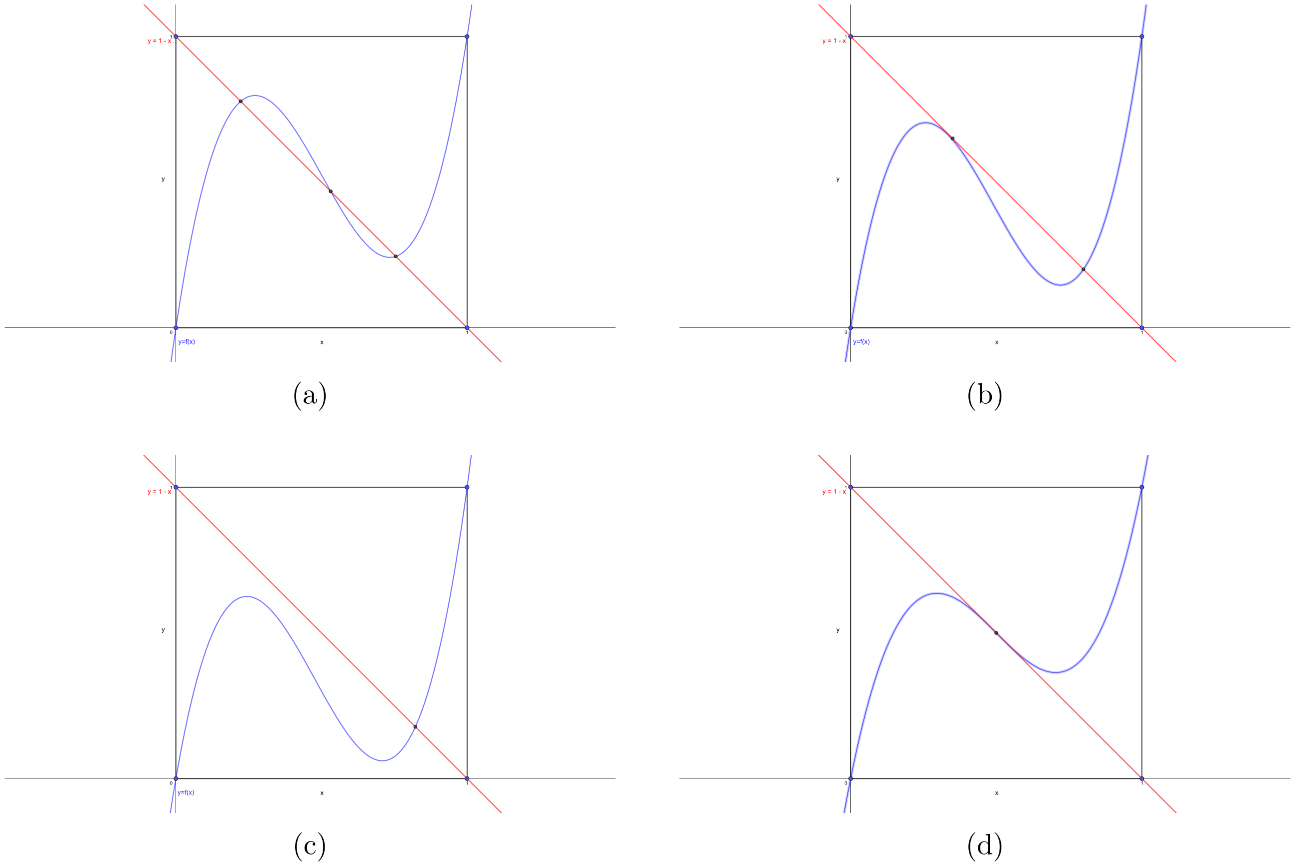


Figure 1: Number of equilibria of System (8) according to the intersection between the line $v = 1 - x$ and the cubic $v = f(x)$ in the unit square.

Note that when the system has three equilibria on [0,1] (as in Figure 1a), the latter are simple roots of the function $g$. When the system has two equilibria on [0,1] (as in Figure 1b), one of them is of multiplicity two. Finally, if the system has only one equilibrium, it can be of single or triple multiplicity (as seen in Figures 1c and 1d, respectively).

As can be seen in Figures 4a and 4b, the local stability of an equilibrium shapes the limiting behavior of the system around that point. The nature of the equilibrium depends on the signs of the two eigenvalues of the Jacobian matrix evaluated at that point. These signs are captured by the signs of the Jacobian matrix's trace and determinant. The trace is the sum of the eigenvalues, while the determinant is their product. Figure 2 summarises the different types of possible equilibria's local stability as a function of the signs of the trace and of the determinant.

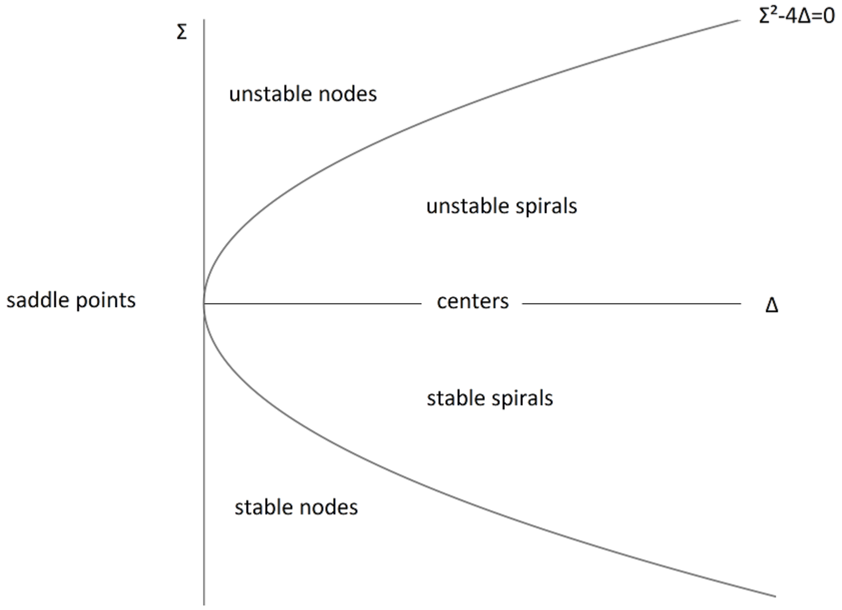


Figure 2: Local stability of an equilibrium according to the sign of the trace Σ and the determinant Δ of the Jacobian matrix.

The Jacobian matrix of System (8) is given by

$$J(x,v) = \begin{pmatrix} -f'(x) & 1 \\ -r(cv+1-c) & -r[(cv-c+1)+c(v+x-1)] \end{pmatrix},$$

where $f'(x) = 3bx^2 + 2(a-b)x + (1-a)$. For $(x^*, v^*)$ a stationary point, we have

$$J(x^*,v^*) = \begin{pmatrix} -f'(x^*) & 1 \\ crx^* - r & crx^* - r \end{pmatrix}.$$

If both eigenvalues of $J(x^*, v^*)$ are positive then the equilibrium point $(x^*, v^*)$ is repulsive. If they both are negative then $(x^*, v^*)$ is attractive and if they have opposite signs; then $(x^*, v^*)$ is a saddle point (since $\Delta < 0$, as seen in Figure 2). Let us thus study the signs of the eigenvalues using trace and determinant. We have

$$\begin{aligned} \operatorname{tr}(J(x^*,v^*)) &= -r(1-cx^*) - f'(x^*), \\ \det(J(x^*,v^*)) &= r(1-cx^*)(f'(x^*)+1). \end{aligned} \tag{11}$$

The signs of the trace and determinant depend on the position of $f'(x)$ relatively to $r(cx^* - 1)$ and $-1$ as indicated in the next proposition. The proof is left to the reader.

**Proposition 3.** *Let $(x^*, v^*)$ be an equilibrium point of System (8).*

- *If $f'(x^*) < -r(1-cx^*)$ and $f'(x^*) > -1$, then $(x^*, v^*)$ is locally stable.*
- *If $f'(x^*) > -r(1-cx^*)$ and $f'(x^*) > -1$, then $(x^*, v^*)$ is locally unstable.*
- *If $f'(x^*) < -1$, then $(x^*, v^*)$ is a saddle point.*

From Proposition 3 in the case of three equilibria, we can infer the local stability of each equilibrium.

**Proposition 4.** *If System* (8) *has three distinct stationary points* $x_1^* < x_2^* < x_3^*$*, then* $x_2^*$ *is a saddle point.*

*Proof.* The assumption allows us to express (10) as

$$f(x) + x - 1 = (x - x_1^*)(x - x_2^*)(x - x_3^*).$$

Thus, $\det(J(x_2^*, v_2^*)) = r(1 - cx_2^*)(f'(x_2^*) + 1) = r(1 - cx_2^*)(x_2^* - x_1^*)(x_2^* - x_3^*) < 0$, which implies that the two eigenvalues have opposite signs and $x_2^*$ is a saddle point. □

The stability information obtained from the eigenvalues gives us the local behavior around the stationary points. Nevertheless, when the equilibrium is locally stable, this does not exclude the existence of a stable limit cycle in the domain $[0, 1]^2$ (see Figure 4c). To exclude the existence of such a cycle, we can either use the Poincaré-Bendixson Theorem [5, Theorem. 1.8.1 p.44] or find a Dulac function [5, Theorem. 1.8.2 p.44]. Unfortunately finding a Dulac function seems out of reach because of the great asymmetry of the system. We have obtained a partial answer in the following specific case.

**Proposition 5.** *If* $-f'(x) - r((cv - c + 1) + c(v + x - 1)) > 0$ *for all* $(x, v) \in [0, 1]^2$*, then there is no stable limit cycle in the domain* $[0, 1]^2$.

*Proof.* The assumption of Proposition 5 implies that on $[0, 1]^2$, the function $|\text{tr}(J(x, v))|$ is different from 0 and keeps the same sign. Using the Bendixson-Dulac Theorem, we can conclude that there is no stable limit cycle in the domain. □

Note that one can find parameters $a, b, c$ or $r$ that do no satisfy the assumption of Proposition 5. In Figure 4c, we exhibit a set of parameters leading to a stable fixed point and a stable limit cycle. These behaviors can be generated through bifurcations, see below.

#### 3.3 Bifurcation analysis

Based on [5] and [6], we study the nature of the bifurcations in System (8). To this aim, we consider System (8) as function of one parameter $\mu$ which can be either $a$, $b$, $c$ or $r$. Thus we rewrite the system as

$$\dot{Z} = G(Z, \mu), \text{ where } Z = \begin{pmatrix} x \\ v \end{pmatrix}, \ \mu \in \mathbb{R} \text{ and } G : \mathbb{R}^2 \times \mathbb{R} \to \mathbb{R}^2. \tag{12}$$

Note that a bifurcation point $(Z^*, \mu^*)$ of System (12) is a set of points for which the flow is structurally unstable. We study two types of bifurcations: saddle-node and Hopf bifurcations. A saddle-node bifurcation is the emergence (or disappearance) of two equilibria, one stable and one unstable when the parameter $\mu$ varies (as in Figure 3a), whereas a Hopf bifurcation changes the local flow around the stationary point (as in Figure 3b).

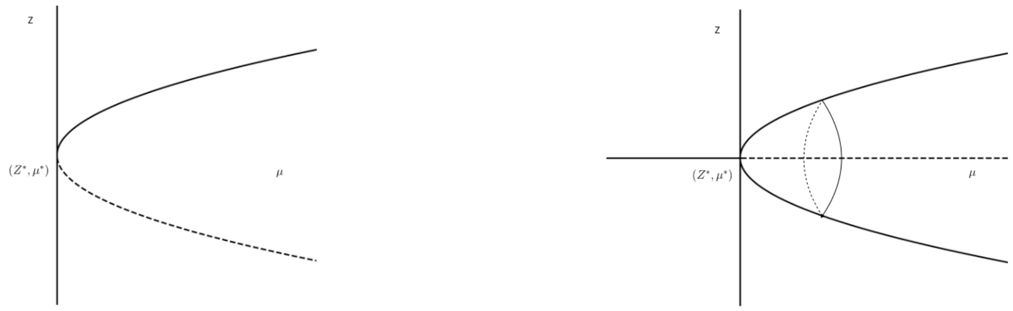


(a) Saddle-node bifurcation (b) Hopf bifurcation

Figure 3: Plain lines are stable stationary points, dashed lines are unstable ones and $(Z^*, \mu^*)$ is a bifurcation point.

From (10) and (9), it is obvious that only $a$ and $b$ can affect the value and the number of stationary points in System (6). Therefore only $\mu = a$ or $\mu = b$ can generate a saddle-node bifurcation. The next proposition gives a simple condition ensuring that these bifurcations are generic.

**Proposition 6.** *Assume that* $(a, b) \neq (-4, 8)$*. Then, the saddle-node bifurcations of System* (12) *are generically non-degenerate for* $\mu = a$ *or* $b$*.*

*Proof.* To characterize the non-degenerate saddle-node bifurcations, we will apply Theorem 3.4.1 from [5, p.148]. When $\mu = \mu^*$, one must check that there exists a stationary point for which

1. $\frac{\partial G}{\partial Z}(Z^*, \mu^*)$ has a simple zero eigenvalue associated to the left eigenvector $w$ and its other eigenvalue has a real part;
2. $w \cdot \frac{\partial G}{\partial \mu}(Z^*, \mu^*) \neq 0$;
3. $w \cdot \frac{\partial^2 G}{\partial Z^2}(Z^*, \mu^*)(v, v) \neq 0$.

The above conditions are satisfied if the stationary point is not a triple multiplicity root of $g$. We find by identification that this is only the case when $(a, b) = (-4, 8)$.

□

From (11), we note that $c$ and $r$ can affect the value of the Jacobian matrix's eigenvalues and thus generate a Hopf bifurcation. Therefore we study the existence of such bifurcations for $\mu = r$ or $\mu = c$ and their non-degeneracy using Theorem 3.4.2 [5, p.151].

**Proposition 7.** *Let* $(Z^*, \mu^*)$ *be the stationary point of System* (12) *with* $Z^* = (x^*, v^*)$*. Assume that* $f'(x^*) < -1$ *and that there exists* $r$*, respectively* $c$*, such that* $r(cx^* - 1) = f(x^*)$*. Then system* (12) *admits a Hopf bifurcation for parameter* $\mu = r$ *respectively* $\mu = c$*, and the bifurcation is generically non-degenerate.*

*Proof.* Under the assumption of Proposition 7, $\det(\frac{\partial G}{\partial Z}(Z^*, \mu^*))$ is non positive and one can find $\mu = r^*$ or $c^*$ such that $\mathrm{tr}(\frac{\partial G}{\partial Z}(Z^*, \mu^*)) = 0$. So, the Jacobian matrix's eigenvalues are pure imaginary and System (12) has a Hopf bifurcation around $(Z^*, \mu^*)$.

We have $\frac{\partial}{\partial r}\mathrm{tr}(\frac{\partial G}{\partial Z}(Z^*, r^*)) = cx^* - 1 \neq 0$ and $\frac{\partial}{\partial c}\mathrm{tr}(\frac{\partial G}{\partial Z}(Z^*, c^*)) = rx^* \neq 0$. Therefore we can apply Theorem 3.4.2 [5, p.151] for $r$ and $c$.

□

##### 3.4 Long-term behaviour of the dynamical system (6)

Figure 4 shows the diversities of stationary states that the dynamic system (6) can admit. Figures 4a, 4b and 4c show the possible behaviors when Eq. (10) vanishes only once, i.e. when the dynamical system admits a unique equilibrium point. Depending on the nature of the equilibrium point, the system can behave quite differently. If this equilibrium point is unstable, then the system admits a stable limit cycle. Figures 4a and 4c show that even if the equilibrium point is stable, this stability is local and we can have the presence of a stable limit cycle .

Figure 4c shows different trajectories and where they converge depending on initial conditions. The black trajectory converges towards the stable equilibrium point while the red and blue trajectories reach the stable limit cycle. The dashed line curve describes the unstable cycle that separates the equilibrium point from the stable cycle.

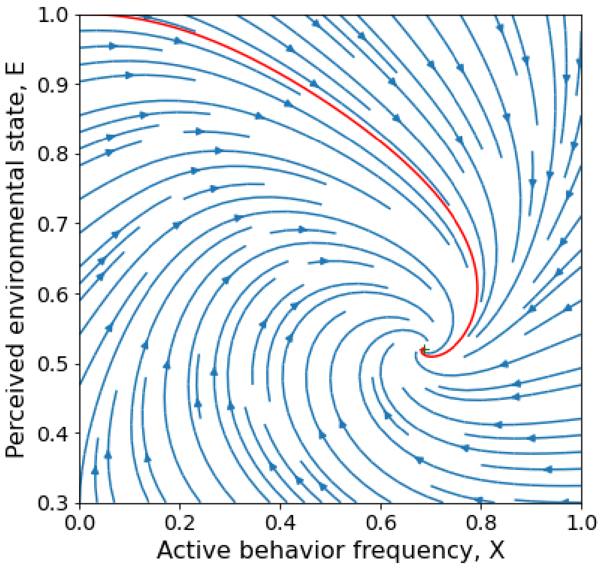


(a) One stable equilibrium

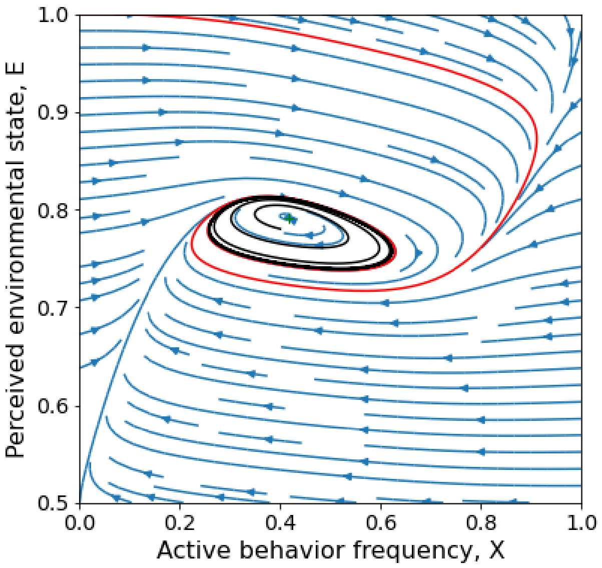


(b) One unstable equilibrium

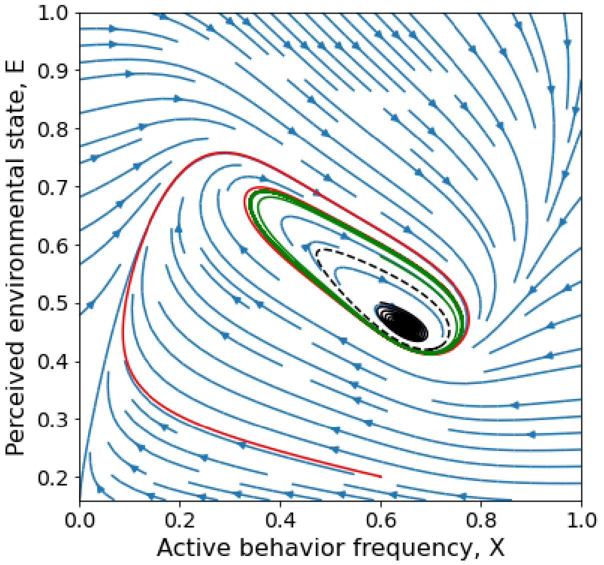


(c) One stable equilibrium with a stable cycle

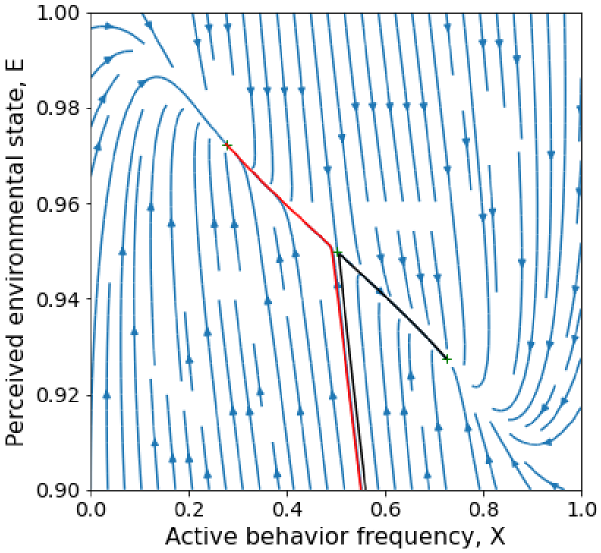


(d) Two stable equilibria and one unstable

Figure 4: Phase portrait of the different behaviors admitted by the dynamic system (6)

Figure 4d shows a situation with two stable and one unstable fixed points, where the steady state depends on the basin of attraction in which the system initially starts.

Overall, three main limit behaviours can be observed: (i) the system admits a stable equilibrium, (ii) the equilibrium is unstable and admits a stable cycle; (iii) two stable equilibria, hence the system is bistable.

### 4 Fluctuations of the stochastic model around its deterministic limit

We study the fluctuations of the stochastic process $(X_t^N, E_t^N)$ around the deterministic function $(x_t, e_t)$ which are defined by the process $\eta_t^N = \sqrt{N}\left(X_t^N - x_t, E_t^N - e_t\right)$.

We rewrite the process $(\eta_t^N)_{t\leq 0}$ as

$$\eta_t^N = \eta_0^N + \int_0^t J^N(X_s^N, E_s^N, x_s, e_s)\eta_s^N + R^N(X_s^N, E_s^N, x_s, e_s)ds + \tilde{M}_t^N, \tag{13}$$

where $J^N(X^N, E^N, x, e)$ is the following matrix

$$J^N(X^N, E^N, x, e) = \begin{pmatrix} J_{11}^N(X^N, E^N, x, e) & J_{12}^N(X^N, E^N, x, e) \\ J_{21}^N(X^N, E^N, x, e) & J_{22}^N(X^N, E^N, x, e) \end{pmatrix}$$

with

$$\begin{aligned}
J_{11}^N(X^N, E^N, x, e) &= \kappa((1 - (X^N + x))N\lambda_{AB}^N(X^N) + (x - x^2)(\delta_A + \delta_B)) + \tau(l_A - l_B),\\
J_{12}^N(X^N, E^N, x, e) &= \tau,\\
J_{21}^N(X^N, E^N, x, e) &= -\ell e(l_B - l_A),\\
J_{22}^N(X^N, E^N, x, e) &= \ell(l_A X^N + l_B(1 - X^N) - (E^N + e))
\end{aligned}$$

and $\lambda_{AB}^N(x) = \lambda_A^N(x) - \lambda_B^N(x)$, $\lambda_{AB}(x) = \lambda_A(x) - \lambda_B(x)$.

Moreover, $R^N(X^N, E^N, x, e) = \begin{pmatrix} \sqrt{N}(x - x^2)(N\lambda_{AB}^N(X^N) - \lambda_{AB}(X^N)) \\ 0 \end{pmatrix}$ and $\tilde{M}^N$ is a square-integrable martingale with quadratic variation given by

$$\langle \tilde{M}^N \rangle_t = \int_0^t \begin{pmatrix} \kappa X_s^N(1 - X_s^N)N[\lambda_A^N(X_s^N) + \lambda_B^N(X_s^N)] + \tau(E_s^N - l_A)(1 - X_s^N) + (l_B - E_s^N)X_s^N \\ 0 \end{pmatrix} ds.$$

Note that the quadratic variation of $\tilde{M}^N$ is of order 1 and will not vanish when N tends to infinity.

We introduce the following assumption: for all $x \in [0, 1]$,

$$\lim_{N\to+\infty} \sqrt{N}(N\lambda_{AB}^N(x) - \lambda_{AB}(x)) = 0. \tag{14}$$

The next theorem describes the limiting behavior of the process $\eta^N$ when N tends to infinity.

**Theorem 8.** *When $N \to \infty$, under Assumption* (14) *and if $(\eta_0^N)_{N\in\mathbb{N}^*}$ converges in law toward $\eta_0$, then for all $T > 0$, the sequence $((\eta_t^N)_{N\in\mathbb{N}}, t \in [0, T])$ converges in law in $\mathbb{D}([0, T], [0, 1] \times [l_A, l_B])$ to the unique solution of:*

$$\eta_t = \eta_0 + \int_0^t J(x_s, e_s)\eta_s ds + \int_0^t D_s dW_s, \tag{15}$$

*where* $(W_t, t \geq 0)$ *is a Brownian motion,* $J(x,e)$ *the matrix given by*

$$J(x,e) = \begin{pmatrix} J_{11}(x,e) & J_{12}(x,e) \\ J_{21}(x,e) & J_{22}(x,e) \end{pmatrix}$$

*with*

$$\begin{aligned}
J_{11}(x,e) &= \kappa((1-2x)(\gamma_A - \gamma_B - \delta_B + (\delta_A + \delta_B)x) + (x - x^2)(\delta_A + \delta_B)) + \tau(l_A - l_B), \\
J_{12}(x,e) &= \tau, \\
J_{21}(x,e) &= -\ell(l_B - l_A)e, \\
J_{22}(x,e) &= \ell(l_A x + l_B(1-x) - 2e)
\end{aligned}$$

*and* $D_s = \begin{pmatrix} \sqrt{\kappa x_s(1-x_s)(\gamma_A + \gamma_B + \delta_A x_s + \delta_B(1-x_s)) + \tau((1-x_s)(e_s - l_A) + x_s(l_B - e_s))} \\ 0 \end{pmatrix}.$

The proof is similar to the one developed in the supplementary materials of [2]-Section 4.

### 5 Simulations and algorithm

We use the following algorithm to simulate the PDMP and overcome the problem with the coupled dynamics of the jumping and continous parts of the process. This algorithm is similar to an acceptance-rejection algorithm. We take the highest jump rate to simulate the time at which the process jumps and we update the current system's state depending on the outcomes. When the jumping time occurs, three events are possible: either an individual switch from behavior A to behavior B, or vice versa, or no switch happens in the population.

**Algorithm 1** Simulation of the stochastic process $(X_t^N, E_t^N)$ in $[0, T_{max}]$

**Initialize the system's state:**
Time: $t \leftarrow t_0$
System's initial state:
Mitigator frequency: $x \leftarrow x_0$
Perceived environment: $e \leftarrow e_0$
Jump rate : $\xi > \sup_{x,e}(N(N\kappa(\lambda_A^N(x) + \lambda_B^N(x)) + \tau_A(e) + \tau_B(e)))$
**while** $t \leq T_{max}$ **do**
  Draw two pseudorandom numbers, $r_1$ and $r_2$ in $[0, 1]$
  Time before next jump: $t_1 \leftarrow -\frac{\log(r_1)}{\xi}$
  Gain probability of an individual in active behavior: $p_+ \leftarrow \frac{N(1-x)(N\kappa x\lambda_A^N(x)+\tau_A(e)))}{\xi}$
  Loss probability of an individual in active behavior: $p_- \leftarrow \frac{Nx(N\kappa(1-x)\lambda_B^N(x)+\tau_B(e))}{\xi}$
  Save and update:
  $t \leftarrow t + t_1$
  $e \leftarrow e + t_1 \times h(x, e)$
  *Three possible outcomes in the population: an individual with baseline behavior becomes active, an individual with active behavior switches to baseline, or no jump*
  **if** $r_2 \leq p_+$ **then**
    *An individual in baseline behavior becomes active:* $x \leftarrow x + \frac{1}{N}$
  **else if** $p_+ < r_2 \leq p_+ + p_-$ **then**
    *An individual in active behavior switches to baseline:* $x \leftarrow x - \frac{1}{N}$
  **else** $\{r_2 > p_+ + p_-\}$
    *No jump from one behavior to the other:* $x \leftarrow x$
  **end if**
**end while**
